# tdLanYFP, a yellow, bright, photostable and pH insensitive fluorescent protein for live cell imaging and FRET-based sensing strategies

**DOI:** 10.1101/2021.04.27.441613

**Authors:** Yasmina Bousmah, Hana Valenta, Giulia Bertolin, Utkarsh Singh, Valérie Nicolas, Hélène Pasquier, Marc Tramier, Fabienne Merola, Marie Erard

## Abstract

Yellow fluorescent proteins (YFP) are widely used as optical reporters in Förster Resonance Energy Transfer (FRET) based biosensors. Although great improvements have been done, the sensitivity of the biosensors is still limited by the low photostability and the poor fluorescence performances of YFPs at acidic pHs. Here, we characterize the yellow fluorescent protein, tdLanYFP, derived from the tetrameric protein from the cephalochordate *B. lanceolatum*, LanYFP. With a quantum yield of 0.92 and an extinction coefficient of 133 000 mol^−1^.L.cm^−1^, it is, to our knowledge, the brightest dimeric fluorescent protein available. Contrasting with EYFP and its derivatives, tdLanYFP has a very high photostability *in vitro* and in live cells. As a consequence, tdLanYFP allows imaging of cellular structures with sub-diffraction resolution using STED nanoscopy and is compatible with the use of spectro-microscopies in single molecule regimes. Its very low pK_1/2_ of 3.9 makes tdLanYFP an excellent tag even at acidic pHs. Finally, we show that tdLanYFP is valuable FRET partner either as donor or acceptor in different biosensing modalities. Altogether, these assets make tdLanYFP a very attractive yellow fluorescent protein for long-term or single-molecule live cell imaging including FRET experiments at acidic pH.

---

Fluorescent proteins (FPs) are used as optical reporters in various biosensing modalities to explore cellular chemistry or cell signaling pathways *in situ*. In particular, FPs are often used in experiments taking advantage of the Förster Resonance Energy Transfer (FRET) phenomenon ^1^. For FRET, a pair of donor/acceptor FPs is required, with the fluorescence spectrum of the donor overlapping the absorption one of the acceptor ^2^. If the donor and the acceptor are fused to two proteins of interest, FRET can be used to probe protein-protein interactions ^3^. It is also possible to fuse them on both sides of a sensing module to build the so-called FRET intramolecular biosensors, which are useful tools to monitor variations in ion or metabolite concentrations ^1^ or to follow enzymatic activities^4^ in living cells. The sensitivity and the reliability of FRET-based biosensors depend on the specifications of the donor/acceptor pair such as a simple photophysical behavior with a mono-exponential fluorescence decay and limited excited-state reactions, an efficient maturation and a high molecular brightness.

In most FRET experiments (~60%), a cyan FP is chosen as the donor and a yellow one as the acceptor^5^,^6^. For CFPs, variants with a low sensitivity to their environment, an efficient maturation, a high brightness including quantum yield close to 1, and a good photostability are now available (Aquamarine ^7^, mTurquoise ^8^, mTurquoise2 ^9^). The best ones notably display a low pK_1/2_, the pH value, at which 50% of the fluorescence intensity is lost, down to 3.1, which is compatible with detection of biological events in the acidic environment ^5^. This feature opens opportunities to develop FRET-based biosensors dedicated to the real-time visualization of metabolic activities in acidic environments, whether it is due to local pH variations or due to their localization in intracellular compartments ^10^, ^11^, ^12^.

Concerning the acceptor, EYFP served as a template for several rounds of engineering leading to improved versions of YFPs as mCitrine ^13,14^ or mVenus ^15^ and Citrine2 ^16^. Citrine, is less sensitive to chloride ions and pH than EYFP but displays a poor photostability ^13^. The improvement of the photostability of mCitrine lead to Citrine2, which additionally retained a high brightness ^16^. Despite these efforts, none of the YFPs derived from EYFP are compatible with the reporting of biochemical processes in an acidic environment so far, their pK_1/2_ ranging from 5.3 to 6 ^16–18^. Consequently, no FRET biosensors based on a cyan/yellow pair were developed for measurements at low pHs ^19^.

LanYFP, a yellow tetrameric FP from *Branchiostoma lanceolatum* has been intensively engineered by Shaner *et al*. leading to the blue-shifted monomeric variant mNeonGreen ^17,20^. While LanYFP has a low pK_1/2_ of 3.5 ^17^, the pH sensitivity of mNeonGreen is equivalent to that of mCitrine (pK_1/2,mNeonGreen_ ~5.4-5.7) ^17,18^. Yet, mNeonGreen has a better photostability than mCitrine under the same irradiation conditions ^17,18^. mNeonGreen’s brightness is also slightly better (~ 1.5 times) than all available yellow variants ^17,18^. Following these observations, we became interested in the intermediate dimeric version, dLanYFP, obtained along the genetic engineering of mNeonGreen.. Here, we present a complete characterization of dLanYFP *in vitro* and evaluate the potential given by its tandem version, tdLanYFP, for live cell imaging. Especially, we highlight its performance as FRET partner in genetically encoded biosensors even at acidic pH, and its suitability for super-resolution microscopy thanks to its very high photostability.

## Results and discussion

### Biochemical parameters and fluorescence characterization of dLanYFP

dLanYFP was produced as a recombinant protein in *E. coli*. Although it is expressed as a monomer, a strong spontaneous dimer formation was observed with size exclusion chromatography and on SDS-PAGE where these dimers dissociated only after heating the sample (Fig. S1). We compared the spectroscopic properties of purified dLanYFP to those of the widely used EYFP, Citrine and Venus. All absorption and fluorescence spectra are equivalent, with a slight blue shift of the absorption and emission maxima of dLanYFP’s spectra by 3 nm and 8 nm respectively (Fig. S2, Table 1). Therefore, dLanYFP can directly replace standard YFPs without adaptation of spectral selections. Its molar extinction coefficient, evaluated per chromophore with the BCA assay, is 133 000 mol^−1^.L.cm^−1^ and its quantum yield 0.92 (Table 1). Those values are consistant with the ones reported by Shaner *et al* ^17^. The relative brightness of the fluorophore in dLanYFP is thus improved by nearly a factor of two compared to EYFP, which is remarkable considering that their chromophores are identical (Gly-Tyr-Gly). dLanYFP, together with its parent LanYFP, thus ranges among the brightest fluorescent proteins known today. In live cells, the strong dimer formation by dLanYFP will double the quantity of labeling fluorophores, leading to a potential 4-fold gain in brightness for fluorescence imaging, as compared to the tagging with conventional YFPs. The fluorescence decays of the purified proteins were acquired using the time-correlated single-photon counting (TCSPC) technique upon excitation at 515 nm (Fig. S3, Table 1). For Venus, Citrine and EYFP, the decays are well fitted by near-single exponentials (> 92-94% of the amplitude), with lifetimes in the range of 3 ns consistent with literature ^21^. The decay of dLanYFP is also mostly mono-exponential (> 97%), with a lifetime in a similar range (2.9±0.1 ns). Interestingly, it also includes a small contribution of a negative, very short component, not present in other YFPs, and whose proportion depends on the emission wavelength, suggesting some fast reaction in the excited-state (Fig. S3).

**Table 1:** *In vitro* characterization of YFPs.

|  | **dLanYFP** | **EYFP** | **Venus**<br>(EYFP F46L-F64L-M153T-V163A-S175G) | **Citrine**<br>(EYFP Q69M) |
| --- | --- | --- | --- | --- |
| $\lambda_{\max}$ excitation (nm) | 512 | 514 | 515 | 515 |
| $\lambda_{\max}$ emission (nm) | 519 | 527 | 528 | 527 |
| Extinction coefficient<br>per chromophore ($\text{mol}^{-1}\text{Lcm}^{-1}$) | $133000 \pm 8000^1$ | $84000 \pm 5000$ | $107000 \pm 6400$ | $91000 \pm 5500$ |
| Quantum Yield | $0.92 \pm 0.05$ | $0.79 \pm 0.05$ | $0.78 \pm 0.05$ | $0.83 \pm 0.05$ |
| Rel. Brightness per chain | 1.84 | 1 | 1.27 | 1.08 |
| Ave. fluo. lifetime<sup>2</sup> ($\pm 0.1$ ns) | $2.9^3$ | 3.2 | 2.9 | 3.3 |
| pK $\frac{1}{2}$ ($\pm 0.1$), $[\text{Cl}^-] = 85 \text{ mM}^4$ | 3.9 | 6.6 | 5.9 | 5.7 |
| Chloride affinity, $K_{\text{app}}$ (mM)<sup>5</sup> | $135 \pm 40$ | $70 \pm 20$ | > 250 | > 300 |
| Photobleaching decay time (s) on<br>agarose beads<sup>6</sup> | $687 \pm 15$ | $102 \pm 20$ | $49 \pm 10$ | $37 \pm 10$ |
| Photobleaching decay time (s)<br>live cells<sup>6</sup> | $463 \pm 15$ | $44 \pm 5$ | ND | $22 \pm 5$ |

1 : for one monomer, 2 : exc 515nm, em 580nm, 20°C, pH8., 3 : neg. exponent 0.19ns, 3% @ em 535nm, <0.5% @ em 580 nm, 4 : values obtained with the processing of the absorption spectra, 5 : affinities are the average of the values derived from the fits of the data obtained from the processing of the absorption and the fluorescence spectra, 6 : under constant wide-field illumination ex 500 nm, em 542 nm, 0.4W/cm<sup>2</sup>

The irreversible photobleaching rate of dLanYFP was compared with the ones of other YFPs immobilized on agarose beads (Fig. S4, Table 1). Under a constant illumination into their major absorption band performed with a wide-field microscope, the fluorescence intensity decrease was slower (15–20 times) for dLanYFP in comparison to the other YFPs. This makes this YFP attractive for imaging applications under intense or prolonged illumination.

The YFP’s chromophore contains a phenol moiety that exists both in protonated and deprotonated form, the latter one being fluorescent. It has been shown that pK_1/2_ depends on chloride ion concentration, [Cl^−^], which can undergo large variations in biological systems. We compared the pK_1/2_ of dLanYFP with the ones of the YFPs for different chloride ion concentrations in the medium (Fig. S5 and S6, Table 1). Consistently with the literature, the pK_1/2_ of EYFP depends strongly on [Cl^−^] concentration and are higher than the ones of Citrine and Venus ^13,17,22^. For dLanYFP, the pK_1/2_ also depends on [Cl^−^] concentration, but is significantly lower regardless its value in comparison to Citrine, leading to a pK_1/2_ of 3.9±0.1 at 85 mM [Cl^−^], an unprecedented value for a YFP, consistent with the low pK_1/2_ of 3.5 of its parent, LanYFP ^17^.

Finally, we also evaluated the maturation kinetics of dLanYFP that was slower than the one of Citrine (Fig. S7), while remaining in the same order of magnitude or faster than the best CFPs used as donor in combination with YFPs^7^. It is likely that the dimeric character of dLanYFP slowed down the maturation rate compared to the monomeric mNeonGreen^21^. Such phenomenon was also reported for DsRed and its dimeric and monomeric derivatives ^23^.

### The tandem version, tdLanYFP: from expression in live cells to applications in spectro-microscopy

The dimerization tendency of dLanYFP might have deleterious consequences in live cell imaging, by forcing non-physiological intermolecular interactions of its fusion constructs. To remove this drawback, we promoted the formation of a stable intramolecular dimer of dLanYFP by concatenating two copies of the gene separated by a sequence coding for a 19 amino-acid linker (Fig. S8) as it was previously done with other existing FP dimers such as dTomato ^23,24^. We verified that the spectroscopic properties of tdLanYFP *in vitro* were similar to the ones of dLanYFP (Fig S2). To evaluate the tendency of tdLanYFP to oligomerize, the organized smooth endoplasmic reticulum (OSER) assay was performed in comparison with mCitrine (Citrine A206K)^25^. The percentage of COS7 cells transfected with CytERM-tdLanYFP presenting whorled structures of endoplasmic reticulum triggered by FP oligomerization was evaluated and typical examples of OSER positive cases for both proteins are shown in Figure S9. For mCitrine, our evaluation of 95% non-OSER situations is consistent with the literature ^18^. For the tandem, the score of 62% non-OSER cases is in the same range as for tdTomato, (57%), and overcomes Citrine (36%) ^18^. We also observed the appropriate localization of tdLanYFP expressed alone in live cells (Fig. 1A), or in fusion with signal peptides targeting the protein to the plasma membrane (Fig. 1B) or the cytoskeleton (Fig. 1C). In addition, we also fused tdLanYFP to the C-terminus of p67^phox^, a cytosolic protein of the NADPH oxidase complex that is responsible for the superoxide anion production in many cell types ^26^. A representative fluorescence lifetime image (FLIM) of COS7 cells expressing p67-tdLanYFP and the corresponding fluorescence decay are presented in Figure 1D and 1E. The average fluorescence lifetime of tdLanYFP fused to p67^phox^ is similar to that of the purified protein recorded *in vitro* (less than 3% of variation) showing little influence of the concatenation of the momoners and of the fusion on this photophysical parameter.

**Figure 1:**
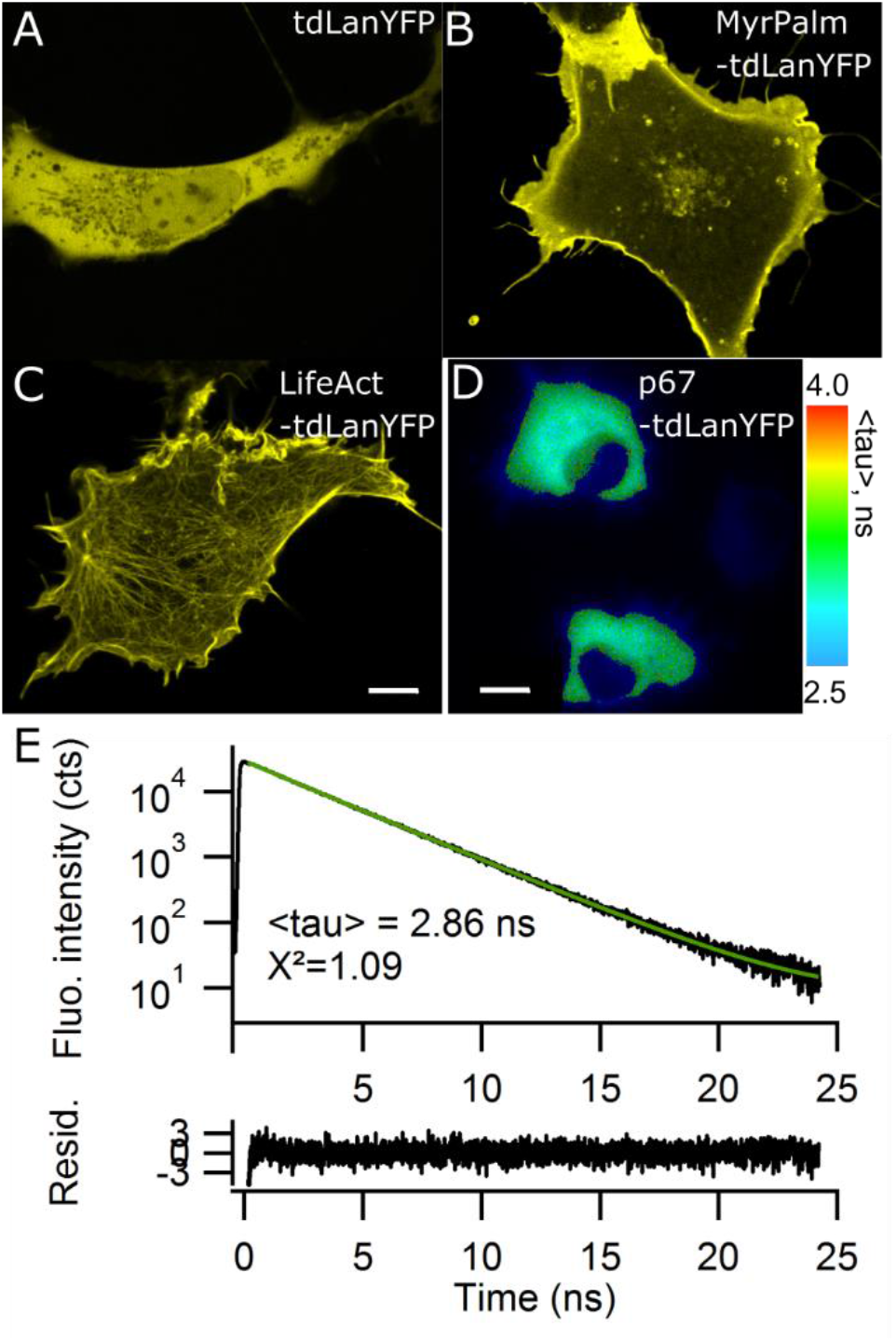
Fluorescence imaging and TCSPC fluorescence decays of tdLanYFP in COS7 cells. Confocal microscopy images of tdLanYFP targeted to the cytosol (A), membrane (B) and cytoskeleton (C). Scale bar 10 μm. (D) Intensity (gray scale) and fluorescence lifetime (color scale) image of a fusion p67-tdLanYFP. Scale bar 15 μm. (E) Fluorescence photons collected from the upper cell were used to calculate the corresponding fluorescence decay. The decay was best fitted with a biexponential fit function (green). The residuals of the fit are displayed below the decay.

The photostability of tdLanYFP was confirmed in live cells, along wide-field and confocal imaging, in comparison to Citrine and EYFP (Fig. S10). Under wide-field illumination, the characteristic time constants recorded in the cell cytosol were close to the one obtained *in vitro* (Table 1). In confocal imaging, FPs were fused to the actin targeting sequence, LifeAct, to avoid any diffusion between the recording of two successive frames. Again, tdLanYFP was more photostable than mCitrine and bleached 2 to 3 times slower (Fig. S10B). The rates of irreversible photobleaching are known to highly depend on the excitation regimes and it is not surprising to observe a difference between (i) homogeneous and continuous wide-field excitation and (ii) the more intense confocal regime where a laser is transiently focused on a single pixel.

Owing to this high photostability, we evaluated the performance of for stimulated-emission-depletion (STED) nano-microscopy using COS7 transiently expressing LifeAct-tdLanYFP (Fig. 2). Due to cell movements between 2 frames, tracking and quantification of the fluorescence intensities of one single filament was subjected to some variability. Nevertheless, we estimated that only 30% of fluorescence was lost after 40 frames. This result gives opportunities for kinetics analysis and organelle tracking with tdLanYFP using STED microscopy.

**Figure 2:**
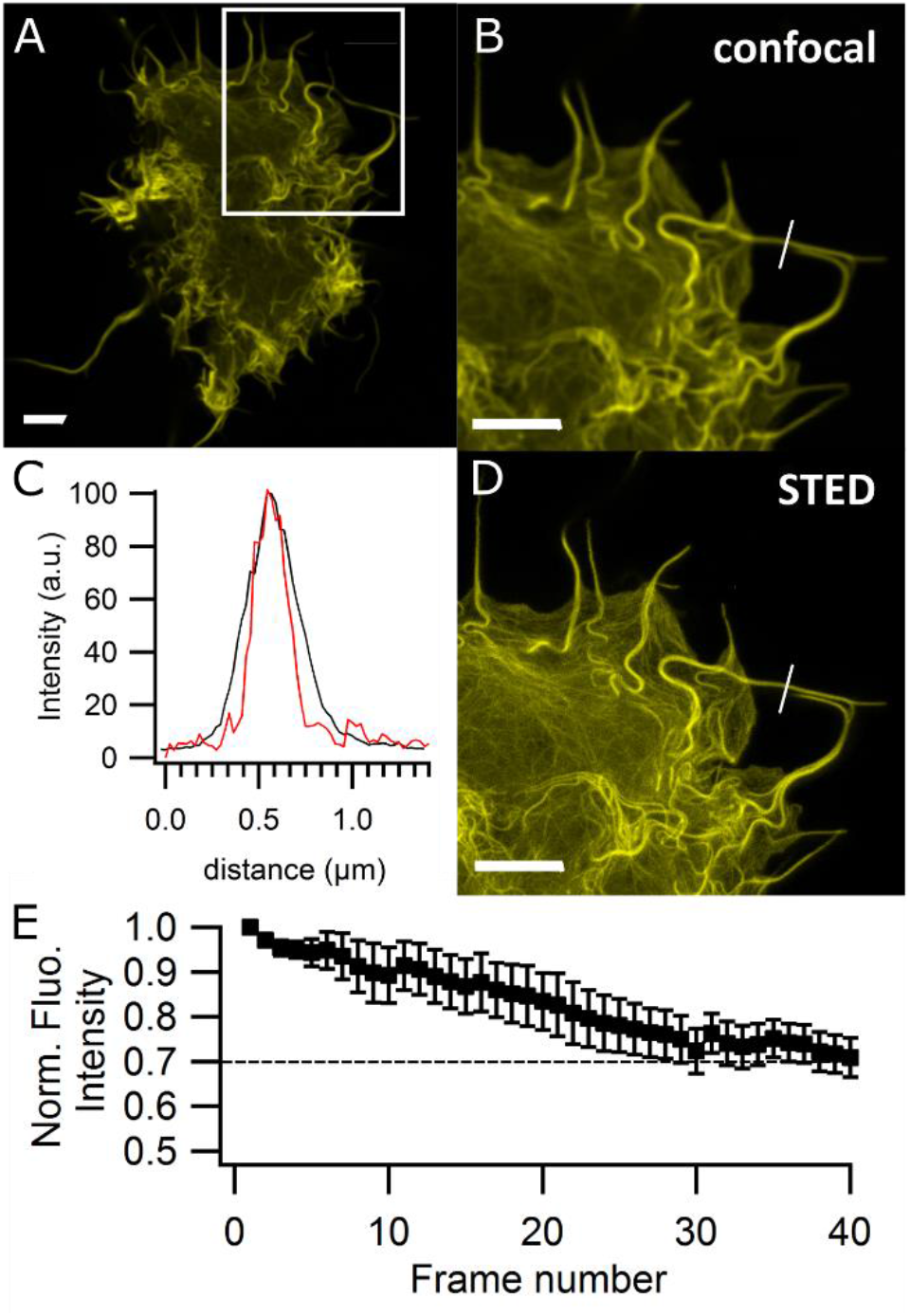
Confocal image of a whole COS7 cell expressing LifeAct-tdLanYFP (A), confocal and STED images of the white ROI (B, D). (C) Graph of the plots profile along the white line to show the gain in resolution. (E) Average of the fluorescence intensity decay in 5 ROIs along the actin filaments of the cell during a 40-frame time-lapse. Scale bars 5 μm.

The combination of high brightness with strong photostability is particularly attractive for experiments in single molecule regimes as for fluorescence correlation spectroscopy (FCS), which monitors the diffusion of a few fluorescent molecules in and out of a confocal volume. In the representative example showed in Figure 3, the fluctuations of the fluorescence intensity of cytosolic tdLanYFP were stable over 60 sec and the amplitude of the autocorrelation function (1/N) of those fluctuations, corresponded to N = 4 fluorophores. This possibility to monitor the fluorescence of only a few molecules is the consequence of the high photostability of tdLanYFP, but it is also due to its improved brightness in comparison to the other YFPs for which there are, indeed, only few examples of live cell FCS experiments despite their widespread use in live-cell fluorescence imaging.

**Figure 3:**
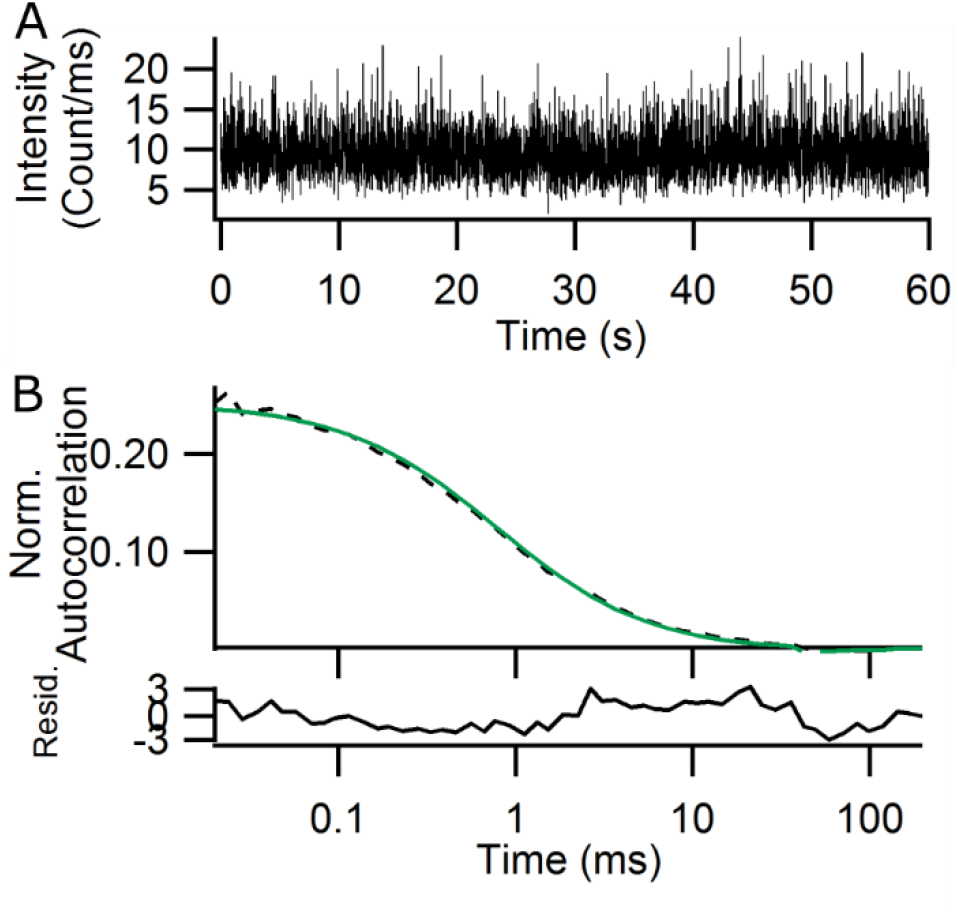
Fluctuations of fluorescence intensity (A) and the corresponding autocorrelation curve (B) acquired in the cytosol of one representative COS7 cell expressing cytosolic tdLanYFP. Experimental values are represented as the dashed line and their fit is in green. The residuals of the fit are displayed below the autocorrelation curve.

### Characterization of a model FRET system: Aquamarine-tdLanYFP FRET fusion

The theoretical Förster distance, R_0_, at which the donor/acceptor FRET efficiency is 50 %, was calculated as 60.4 Å and 67.8 Å for the FRET pair Aquamarine/dLanYFP as monomer or dimer respectively (κ^2^ = 2/3, n=1.33). It is slighty above other CFP – YFP FRET pair (50-60 Å, www.fpbase.org^27^). To evaluate the performance of tdLanYFP as acceptor in FRET-based imaging applications, we built a FRET fusion Aquamarine-tdLanYFP, and compared its response monitored by FLIM in COS7 cells to the previously studied Aquamarine-Citrine fusion (Fig. 4) ^19^. We also recorded the overall fluorescence spectra upon donor excitation in a field of view under the microscope (Fig. S11). The average lifetime of Aquamarine in the Aquamarine-tdLanYFP FRET fusion is significantly lower than the one of Aquamarine expressed alone showing a strong energy transfer from Aquamarine to tdLanYFP, giving an average FRET efficiency of 28 % (Fig. 4). This FRET is also clearly observed in the fluorescence spectrum through the presence of an intense fluorescence band between 520 and 530 nm characteristic of the acceptor fluorescence emission triggered by donor excitation and energy transfer (Fig. S11). The calculated FRET efficiency is similar to the one observed in the original Aquamarine-Citrine construct (32%). This is surprising, considering that in the new construct, each Aquamarine donor is linked to two dLanYFP moieties, each with an increased extinction coefficient in comparison to Citrine. This might arise from unfavorable relative orientation and/or distance of donor and acceptor chromophores in this particular FRET fusion. Alternatively, it might reflect the occurrence of dark, non-absorbing dLanYFP molecules, for example due to incomplete chromophore maturation.

**Figure 4:**
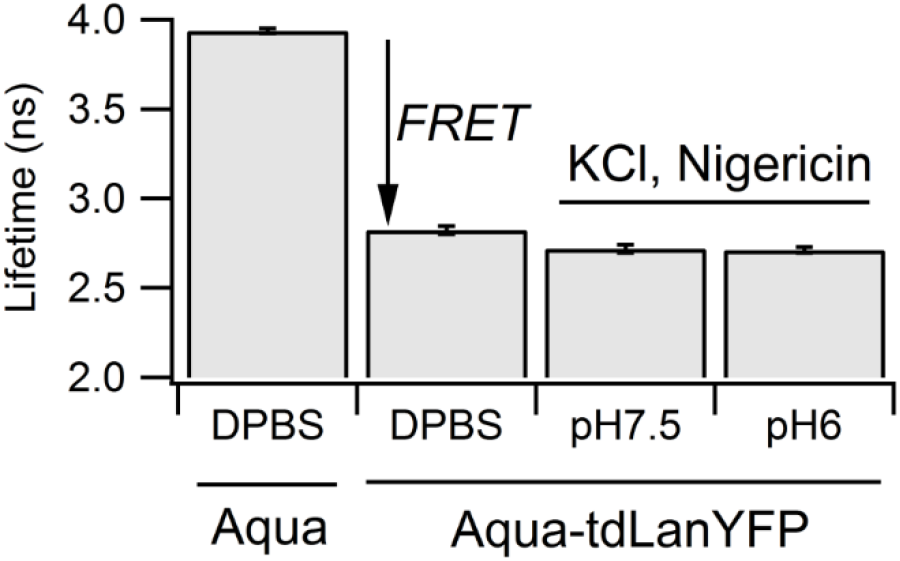
Fluorescence lifetimes of Aquamarine expressed in the cell cytosol of COS7 alone or in tandem with tdLanYFP at different pHs. Average of minimum 20 cells. Values displayed +/−SE.

To evaluate the effect of pH on the FRET efficiency in the construct Aquamarine-tdLanYFP, the intra-cellular pH was fixed at either pH 7.5 or pH 6 using a mixture of MES (15 mM) and HEPES (15 mM) buffers containing 140 mM KCl and 10 μM of nigericin. Within this pH range, Aquamarine’s lifetime did not vary ^7^ and indeed, we observed the same lifetime for FRET fusion at both pHs (Fig. 4). This is an improvement compared to the Aquamarine / Citrine pair whose FRET efficiency changes in function of pH below pH 7 ^19^. Indeed, acidic pHs induce the formation of the protonated form of Citrine whose absorption band, appearing at lower wavelenghts than the deprotonated form, leads to a poor spectral overlay with the donor fluorescence emission spectrum ^19^. As a consequence, the FRET pair Aquamarine / tdLanYFP is thus suitable for FRET studies at acidic pHs at least down to pH 6.

### Applications of tdLanYFP as a FRET partner for live cell imaging

To validate further the Aquamarine / tdLanYFP as a FRET pair, we built a biosensor reporting the autophosphorylation of protein kinase AURKA, tdLanYFP-AURKA-Aquamarine, as well as a kinase-dead version, the mutant AURKA(K162M), as control (Fig. 5A and 5B)^28^. We evaluated their responses by FRET/FLIM on the mitotic spindle in U2OS cells synchronized at mitosis. Aquamarine-AURKA served as lifetime reference. In the case of the AURKA kinase-responsive biosensor version, we observed a lifetime variation of ~ 300 ps corresponding to the FRET from Aquamarine to tdLanYFP upon autophosphorylation. This was a variation slightly higher than the value observed recently with the mTurquoise2/tdLanYFP FRET pair ^28^. We also observed that the kinase-dead version did not completely abolish FRET, neither did the ATP analogue, MLN8237, used to inhibit AURKA autophosphorylation,. This was consistent with previous observations done for the biosensor built with the mTurquoise2/tdLanYFP pair ^28^. As a rule of thumb, an efficient energy transfer can be observed up to twice R_0_, the Förster distance. For Aquamarine/tdLanYFP and mTurquoise2/tdLanYFP FRET pairs, R_0_ are around ~60 Å. For the AURKA kinase-responsive and kinase-dead version of the biosensor, the donor-acceptor distances were estimated to 83 Å and 97 Å respectively ^28^. These two values are below twice R_0_ (~120 Å) and therefore, FRET may be observed even within the more elongated inhibited or kinase-dead versions of the biosensor.

**Figure 5:**
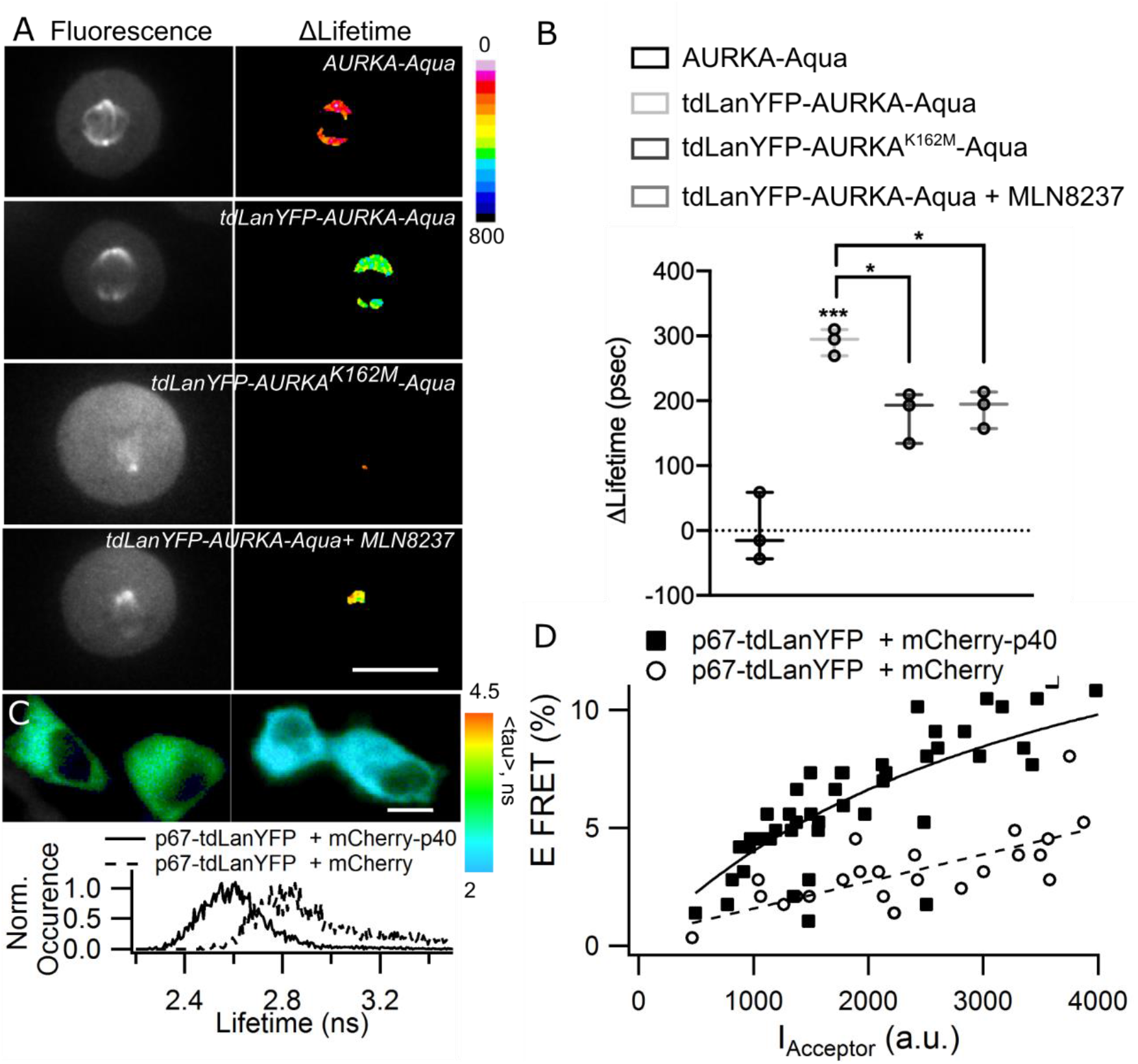
Representative fluorescence intensity (Aquamarine channel) and ΔLifetime images of U2OS cells expressing the indicated constructs and synchronized at mitosis (Scale bar 10 μm) (A) together with the corresponding ΔLifetime quantification at the mitotic spindle (B). The bar in boxplots represents the median; whiskers extend from the 10th to the 90th percentiles. Circles represent the mean FRET values issued by three independent experiments, where 10 cells per experiment were analyzed. (C) Representative FLIM images of COS7 cells expressing p67-tdLanYFP with mCherry (left) or mCherry-p40 (right) and the corresponding lifetime histograms. Scale bar 10 μm. (D) Average FRET efficiencies plotted against acceptor (mCherry) fluorescence intensity. Each symbol represents the values for one cell. Solid and dashed lines are for eye guidance only.

We also evaluated the possibility to use tdLanYFP as a FRET donor in intermolecular FRET. As a model, we used two subunits of the NADPH oxidase, p67^phox^ and p40^phox^ that are known to interact in the resting state of the enzyme ^3^. The subunits were transiently expressed in COS7 cells and FRET was monitored by TCSPC-FLIM (Fig. 5C and 5D). As a reference, we monitored the lifetime of p67-tdLanYFP co-expressed with mCherry. We observed a significant decrease of the donor lifetime when the two protein partners, p67-tdLanYFP and mCherry-p40, were coexpressed (Fig 5C). For each cell, the FRET efficiency was plotted against I_Acceptor_ (Fig. 5D). Interestingly, we noticed a non-specific FRET likely due to crowding that started even at the lowest analysed fluorescence intensities of the acceptor whether the FRET acceptor mCherry is fused to the p40^phox^ subunit or not (Fig. 5D). However, the FRET efficiency for the co-expression of p67-tdLanYFP with mCherry-p40 was always above the control reaching a value of around 10 % for the highest fluorescence intensities of the acceptor consistantly with our previous observations with the same proteins of interest and Citrine as a donor ^3^. It is thus possible to exchange Citrine for tdLanYFP in such experiments.

The use of a dimer as an acceptor may have various consequences on the efficincy of the FRET phenomenon. To follow protein-protein interaction, a more bulky dimer may impair the interaction and appear as a drawback. Nevertheless, the dimer is composed of two potential acceptors with different orientations, which may be beneficial for the energy transfer. Thus, the FRET efficiency, induced by the protein-protein interaction, might be less orientation dependent. For FRET-based biosensors, the benefit may depend on the nature of the conformational change induced by the sensing central module between the donor/acceptor FRET pair. If the conformational change induces a large *distance effect* between completely close and open conformations for example, the use of a dimer may be useful to limit orientational effects. On the contrary, if the FRET variation in the biosensor is mostly related to an *orientation effect* itself between an orientation with minimal FRET in one conformation and maximal FRET in the other, a large FRET variation might be harder to achieve with a dimer, each monomer being a potential efficient acceptor in both conformations.

Finaly, here, we report a lifetime-based FRET detection for the biosensors. For ratiometric detection of FRET on a dedicated microscopy setup, the blue-shifted fluorescence emission spectrum of tdLanYFP in comparison to classical YFPs may necessitate the use of a narrower bandpass emission filter for the detection channel of the donor alone. As a result, this may slightly decrease the signal-to-noise ratio in such ratiometric detection.

## Conclusion

In this work, we show that tdLanYFP is a high-performance yellow fluorescent protein for monitoring intracellular biological processes in various biosensing modalities, as a classic label or to be used in FRET-based strategies. Its characteristics surpass those of the currently available yellow fluorescent proteins derived from the original EYFP.

*In vitro*, dLanYFP keeps most of the oustanding properties of the wild type LanYFP, which are the best features for a yellow fluorescent protein including a brightness per chain 1.7 times higher than Citrine and a fluorescence lifetime of 2.9 ns. The tandem dimeric version, tdLanYFP, retains the same photophysical characteristics as dLanYFP. Constructed for live-cell experiments, it allows most of the conventional imaging applications using the spectral selection available for conventional YFPs. Its photostability is a great advantage for applications requiring high-power illumination or long time observations of a small pool of proteins, for example single molecule tracking or analysis of kinetics even in super-resolved imaging as STED microscopy. Its low pK_1/2_ allows monitoring FRET at pH as low as 6 without any variation of FRET efficiency. This properties pave the way for new applications in acidic environements including the developpement of biosensors for the acidic degradative compartments of the endocytic and autophagic pathways or for the phagosomes, acidic traps of immune cells to destroy microbes. Those machineries are involved in many physio-pathological processes and moleculars tools such as genetically encoded biosensors are still missing mainly due to the lack of reliable FPs able to be a reliable optical reporter at acidic pHs ^12,29^. tdLanYFP was successfuly integrated in a FRET-based biosensor in conjunction with a cyan donor. In most cases, the structural rationale behind a FRET-biosensor, *ie*. the range of conformations probed by the FRET pair involving a whole set of possible distances and orientations, is not known and requires a trial-and-error approach.. In this report, we presented tdLanYFP that will be a valuable probe for any experiment requiring photoastable acceptor with low pH-sensitivity. Even though the tandem can now be used as it is in FRET-based applications, its relatively slow maturation and residual oligomerization tendency could remain a drawback in particular for quantitative hetero-FRET applications. The maturation may be further improved by several rounds of mutagenesis. Nevertheless, the screening for fast-maturating versions of tdLanYFP should be performed while preserving the incredible brightness, excellent photostability and low pH-sensitivity of the current version of the protein.

## Experimental part

Details on molecular biology and *in vitro* experiments can be found in SI.

### Live-cell experiments

Most of the live-cell experiments were performed in COS7. Cells were cultivated in DMEM Glutamax (Invitrogen) supplemented with 5 % FCS in 25 cm^2^ flasks. For microscopy, cells were grown to 80 % confluence on glass coverslip (ibidi) and transiently transfected with the appropriate expression plasmids with XtremeGene HP (Roche Diagnostics) following the supplier’s instructions and used 24–48 h after the transfection.^3^ AURKA FRET biosensor was expressed in U2OS cells. The cells were cultivated, transfected and synchronized at mitosis as previously described. ^28^

### Microscopy

**Wide field bleaching experiments** were performed either on purified 6xHis-tagged fluorescent proteins attached to Ni-NTA agarose beads, or on COS7 cells expressing cytosolic constructs. In both cases, the epifluorescence pathway of the TCSPC-FLIM setup (see below) was used for illumination and fluorescence was excited and collected through a Semrock Brightline cube YFP-2427B-NTE-ZERO. The incident power on the sample was 530 μW and the diameter of the illumination field 400 μm, giving an approximate irradiance of 0.4W/cm^2^. Averages of 5 to 10 individual intensity decay traces were collected for each sample type and fitted with single or double exponential decay models as required, from which the average bleaching time was computed.

**For OSER assay**, COS7 cells expressing the OSER construct of either tdLanYFP or Citrine targeted to the endoplasmic reticulum were imaged with an inverted Leica DMI8 microscope equipped with a 40x/0.6 NA objective (Leica) and piloted with Metamorph software. The epifluorescence pathway was equipped with a solid state light engine (Lumencor), a set of filter cubes and a CCD camera (Flash4.0LT, Hamamatsu Photonics). We used the Semrock Brightline cube YFP-2427B-NTE-ZERO for spectral selection. Images were analyzed visually using ImageJ. Typical examples of OSER cases are shown in Figure S9 for tdLanYFP (750 cells classified) and Citrine (420 cells classified).

**Confocal images** were acquired with an inverted Leica DMI 6000 microscope equipped with a 63x/1.4 NA PL APO oil immersion objective (Leica) and pilot with the LAS-X software (Leica). Excitation was performed at 510 nm with a white light laser and the fluorescence detected between 525 nm and 570 nm with a GaAsP Hybride detector (Hamamatsu).

**For the photobleaching in confocal conditions and for STED imaging**, LifeAct-mCitrine and LifeAct-tdLanYFP expressed in COS7 cells were imaged with an inverted Leica TCS SP8 gated-STED super-resolution microscope (Leica) equipped with a 100x/1.40 NA HCX PL APO STED oil immersion objective and piloted with the Leica SP8 LAS AF software (Version 3.6; Leica). The instrument was equipped with a WLL Laser (511 nm excitation wavelength) and a 592 nm depletion laser. The depletion laser was operated at an average power of 70 % of 400 mW with a gated detection of Tg = 1.5 ns-6 ns. The variations of the fluorescence intensities within the cells during the time-lapse over 30 frames in confocal imaging were measured in several ROIs in the cells and averaged. The final data are the average of 3 different cells.

**For FCS experiments**, an inverted Leica confocal microscope TCS SP8 SMD (Leica) was used. It was equipped with a DMI 6000 CS stand and a 63x/1.2 NA HC PLAN APO water immersion objective. The samples were excited with a continuous argon laser at 514 nm and the fluorescence was selected by 505 nm dichroic mirror and a bandpass filter BP 540/30 nm with an APD detector (PicoQuant). For the detection, the SMD module was constituted of a PicoHarp 3000 system for TTTR mode of single photon counting (PicoQuant). The fluctuations of fluorescence intensity were auto-correlated, and the resulting curves were analyzed using a standard pure diffusion model with SymphoTime (PicoQuant).

**Fast-FLIM** measurements were performed on a Leica DMI6000 microscope (Leica) equipped 63x/1.4 NA oil immersion objective and a time-gated custom-built setup for FLIM ^30^ and driven by the Inscoper hardware (Inscoper). Briefly, cells were excited at 440 ± 10 nm a using a white light pulsed laser (Fianium), and emission was selected using a band-pass filter of 483/35 nm. The detection is time-gated and the fluorescence lifetime was calculated with five sequential temporal gates of 2.2 ns each. The mean pixel-by-pixel lifetime was calculated only when the fluorescence intensity was above 3000 gray values using the Inscoper software (Inscoper). To calculate the ΔLifetime, the mean lifetime of the cells expressing the donor alone (AURKA-Aquamarine) was calculated and its value was used to normalize data in all the analyzed conditions for each experiment.

**TCSPC-FLIM microscopy** was performed using a custom made time-resolved laser scanning TCSPC microscope described previously.^7^ Briefly, the setup is based on a TE2000 microscope with a 60x, 1.2NA water immersion objective (Nikon). The epifluorescence pathway was equipped with a solid state light engine (Lumencor), a set of filter cubes and a CCD camera (ORCA AG, Hamamatsu Photonics). The spectral selections for CFP, YFP and RFP are available Table S1. The CFP cube was used for Aquamarine, the YFP cube for EYFP, Citrine, dLanYFP and tdLanYFP, the RFP cube for mCherry. A spectrometer (Ocean Optics) was fixed on the front exit of the microscope with an optical fiber. Emission spectra were recorded with the CFP excitation filter on the whole field of view (Fig. S11). The TCSPC path was equipped with a 440 and a 466 nm pulsed laser diode (PicoQuant) driven by a PDL800 driver (20 MHz, PicoQuant) for CFP and YFP respectively. The fluorescence was selected by a set of filters (Table S2) before the MCP-PMT detector (Hamamatsu Photonics). The C1 scanning head (Nikon) probed a 100μm x100μm field of view. The PicoHarp300 TCSPC module (PicoQuant) collected all the signals and data were processed with the SymPhoTime64 software (PicoQuant). The TCSPC fluorescence decay of all the pixels of the cytosol was computed by the SymPhoTime64 and decays were fitted with exponential fit functions:

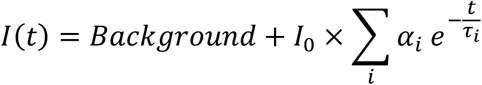

The quality of the fit was evaluated by the weighted residual distribution and Pearson’s χ^2^ test. The average lifetime was computed as < *τ* >= ∑_*i*_ *α*_*i*_ *τ*_*i*_ and the apparent FRET efficiency was calculated using < *τ*_*DA*_ >, the average lifetime of the donor in presence of acceptor and *τ*_*Donor*_, the lifetime of the donor alone:

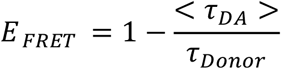

We observed a slight decrease of the donor lifetime when the fluorescence intensity of the acceptor, I_Acceptor_, in the cell increased (Fig. S12). I_Acceptor_ is directly proportional to the concentration of the tandems in the cell cytosol in the case of Figure S12. When their concentration increased, a hetero-molecular FRET between close tandem molecules can happen in addition to the intra-molecular one within the tandems leading to the observed concentration-dependent lifetime decrease^31^. In order to minimize the effect of the so-called molecular crowding on the intra-molecular FRET efficiency, we set a maximum limit for I_Acceptor_ at which the donor lifetime decreased less than 5 % (I_Acceptor_ < 1000 a.u.) in the tandem experiment. In Figure 5, the non-specific FRET due to crowding was directly evaluated by comparing the evolution of FRET efficiencies in presence of mCherry only with the ones in presence of mCherry fused to p40, the interaction partner of p67.

## Supporting information

Supplementary Information_RevisedVersion

## Acknowledgement

This work has benefited from the SpICy microscopy facility of ICP for TSCPC-FLIM measurements, the Microscopy Rennes Imaging Centre (MRic) for FCS and FastFLIM measurements, the Plateforme d’Imagerie Cellulaire MIPSIT for STED measurements and the Light Microcopy Facility Imagerie-Gif for confocal microscopy. We thank P. Pernot for the analysis of the fluorescence decays.

---

For Table of Content Only

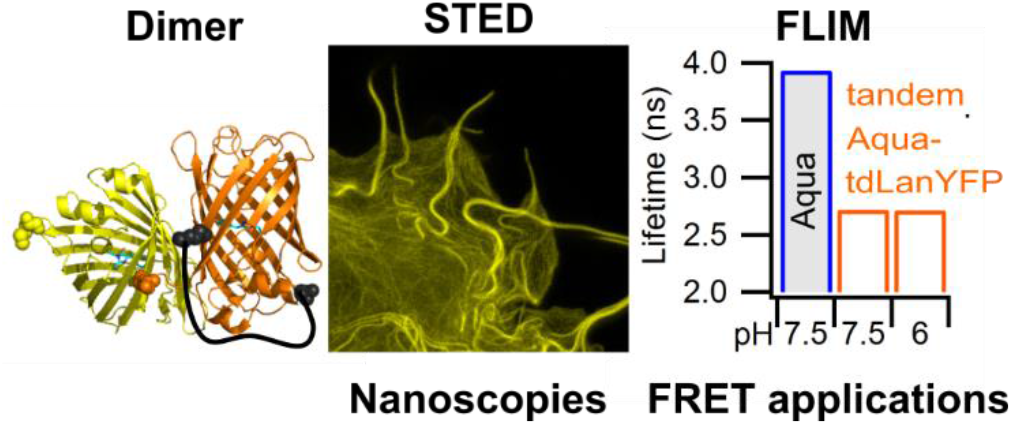

