## Supplementary Information_RevisedVersion for "tdLanYFP, a yellow, bright, photostable and pH insensitive fluorescent protein for live cell imaging and FRET-based sensing strategies"

Molecular Biology and Plasmids p2

In vitro characterization of the proteins p4

Supplementary Figures:

Figure S1 SDS-PAGE and gel filtration chromatography p 7

Figure S2 Normalized absorption and emission spectra of purified yellow fluorescent proteins p 8

Figure S3 Fluorescence decays of purified yellow fluorescent proteins p 9

Figure S4 Irreversible photobleaching of YFPs under continuous irradiation p 10

Figure S5 pH dependence of the fluorescence intensity of YFPs p 10

Figure S6 Chromophore protonation $pK_{1/2}$ of YFPs as a function of chloride concentration p 11

Figure S7 Comparison of the maturation rates of dLanYFP and Citrine p 12

Figure S8 Amino-acid sequence and schematic structure of the tandem tdLanYFP p 13

Figure S9 Wide-field fluorescence images of representative OSER positive cells p 14

Figure S10 Irreversible photobleaching of YFPs in live COS7 cells p 15

Figure S11 Fluorescence spectra of COS7 cells transfected with tandems p 16

Figure S12 Correlation between the donor lifetime and the fluorescence intensity of the acceptor p 16

Supplementary Tables S1 and S2 p 17

References p 18

<u>Molecular Biology and Plasmids</u>

*For in vitro experiments:*

For purification of recombinant 6xHis-tag dLanYFP, the plasmid pNCS-dLanYFP has been brought to Allele Biotechnology and Pharmaceuticals (cat#: ABP-FP-DYPNCS). EYFP, Citrine and tdLanYFP (-***shuffled***) were cloned in pProExHTa [1] and Venus in pBAD (Plasmid #54859, Addgene) to produce a His-tagged protein.

The sequences of the p6xHis-tag purified tdLanYFP (pink) and dLanYFP (black) are:

```
MSYYHHHHHHDYDIPTTENLYFQGA          MVSKGEEDNMASLPATHELHIFGSF
MRGSHHHHHHGMASMTGGQQMGRDLYDDDDKDPMVSKGEEDNMASLPATHELHIFGSF
             NGVDFDMVGRGTGNPNDGYEELNLKSTKGDLQFSPWILVPQIGYGFHQYL
             NGVDFDMVGRGTGNPNDGYEELNLKSTKGDLQFSPWILVPQIGYGFHQYL
             PFPDGMSPFQAAMKDGSGYQVHRTMQFEDGASLTSNYRYTYEGSHIKGEF
             PFPDGMSPFQAAMKDGSGYQVHRTMQFEDGASLTSNYRYTYEGSHIKGEF
             QVKGTGFPADGPVMTNSLTAADWCVTKMLYPNDKTIISTFDWTYTTGNGK
             QVKGTGFPADGPVMTNSLTAADWCVTKMLYPNDKTIISTFDWTYTTGNGK
             RYQSTARTTYTFAKPMAANILKNQPMFVFRKTELKHSKTELNFKEWQKAF
             RYQSTARTTYTFAKPMAANILKNQPMFVFRKTELKHSKTELNFKEWQKAF
             TDVMSTGTGSTGSGSSGEEDNMASLPATHELHIFGSFNGVDFDMVGRGTG
             TDVMGMDELYK MVSKGEEDNMASLPATHELHIFGSFNGVDFDMVGRGTG
             NPNDGYEELNLKSTKGDLQFSPWILVPQIGYGFHQYLPFPDGMSPFQAAM
             NPNDGYEELNLKSTKGDLQFSPWILVPQIGYGFHQYLPFPDGMSPFQAAM
             KDGSGYQVHRTMQFEDGASLTSNYRYTYEGSHIKGEFQVKGTGFPADGPV
             KDGSGYQVHRTMQFEDGASLTSNYRYTYEGSHIKGEFQVKGTGFPADGPV
             MTNSLTAADWCVTKMLYPNDKTIISTFDWTYTTGNGKRYQSTARTTYTFA
             MTNSLTAADWCVTKMLYPNDKTIISTFDWTYTTGNGKRYQSTARTTYTFA
             KPMAANILKNQPMFVFRKTELKHSKTELNFKEWQKAFTDVMGMDELYK
             KPMAANILKNQPMFVFRKTELKHSKTELNFKEWQKAFTDVMGMDELYK
```

*For live cell experiments:*

**p-tdLanYFP-N1** plasmid has been built using a synthetic gene coding for one monomer of tdLanYFP (Eurogentec). Both monomers were cloned successively by PCR in pcDNA3 as an intermediate vector. The gene coding for the tandem was then cloned between AgeI and BsrGI in p-Aquamarine-N1 used as a vector [2]. During the process, a mutation has been introduced to remove BsrGI cloning site in the first monomer.

The final DNA sequence between AgeI and BsrGI (in blue) restriction sites is:

```
 651                       ACCGG TCGCCACCAT GGTCTCCAAA GGAGAGGAGG
 701 ATAACATGGC CTCTCTCCCA GCGACACATG AGTTACACAT CTTTGGCTCC
 751 TTCAACGGTG TGGACTTTGA CATGGTGGGT CGTGGCACCG GCAATCCAAA
 801 TGATGGTTAT GAGGAGTTAA ACCTGAAGTC CACCAAGGGT GACCTCCAGT
 851 TCTCCCCCTG GATTCTGGTC CCTCAAATCG GGTATGGATT CCATCAGTAC
 901 CTGCCCTTCC CCGACGGGAT GTCGCCTTTC CAGGCCGCCA TGAAAGATGG
 951 CTCCGGATAC CAAGTCCATC GCACAATGCA GTTTGAAGAC GGTGCCTCCC
1001 TGACTTCCAA CTACCGCTAC ACCTACGAGG GAAGCCACAT CAAAGGAGAG
1051 TTTCAGGTGA AGGGGACTGG TTTCCCTGCT GACGGTCCTG TGATGACCAA
1101 CTCGCTGACC GCTGCGGACT GGTGCGTGAC CAAGATGCTG TACCCCAACG
1151 ACAAAACCAT CATCAGCACC TTTGACTGGA CTTACACCAC TGGAAATGGC
1201 AAGCGCTACC AGAGCACTGC GCGGACCACC TACACCTTTG CCAAGCCAAT
1251 GGCGGCCAAC ATCCTGAAGA ACCAGCCGAT GTTCGTGTTC CGTAAGACGG
1301 AACTCAAGCA CTCCAAGACC GAACTCAACT TCAAGGAGTG GCAAAAGGCA
1351 TTTACCGATG TGATGTCGAC TGGCACTGGT TCTACGGGCT CGGGCTCCTC
```

```
1401 AGGAGAGGAG GATAACATGG CCTCTCTCCC AGCGACACAT GAGTTACACA
1451 TCTTTGGCTC CTTCAACGGT GTGGACTTTG ACATGGTGGG TCGTGGCACC
1501 GGCAATCCAA ATGATGGTTA TGAGGAGTTA AACCTGAAGT CCACCAAGGG
1551 TGACCTCCAG TTCTCCCCCT GGATTCTGGT CCCTCAAATC GGGTATGGCT
1601 TCCATCAGTA CCTGCCCTTC CCCGACGGGA TGTCGCCTTT CCAGGCCGCC
1651 ATGAAAGATG GCTCCGGATA CCAAGTCCAT CGCACAATGC AGTTTGAAGA
1701 CGGTGCCTCC CTGACTTCCA ACTACCGCTA CACCTACGAG GGAAGCCACA
1751 TCAAAGGAGA GTTTCAGGTG AAGGGGACTG GTTTCCCTGC TGACGGTCCT
1801 GTGATGACCA ACTCGCTGAC CGCTGCGGAC TGGTGCGTGA CCAAGATGCT
1851 GTACCCCAAC GACAAAACCA TCATCAGCAC CTTTGACTGG ACTTACACCA
1901 CTGGAAATGG CAAGCGCTAC CAGAGCACTG CGCGGACCAC CTACACCTTT
1951 GCCAAGCCAA TGGCGGCCAA CATCCTGAAG AACCAGCCGA TGTTCGTGTT
2001 CCGTAAGACG GAACTCAAGC ACTCCAAGAC CGAACTCAAC TTCAAGGAGT
2051 GGCAAAAGGC ATTTACCGAT GTGATGGGCA TGGACGAGC T GTACAAG
```

The repeat of twice the same DNA sequence in the plasmids can complicate further cloning steps. Another version of p-tdLanYFP-N1 was also built with an optimized codon version for human expression (Eurofins). In this construct, the DNA sequence of each monomer is different. When used, this version is called ***shuffled*** in the following.

The final DNA sequence of the ***shuffled*** version between AgeI and BsrGI (in blue) restriction sites is:

```
 651                       ACCGG TCGGCGCCAT GGTCTCCAAA GGAGAGGAGG
 701 ATAACATGGC CTCTCTCCCA GCGACACATG AGTTACACAT CTTTGGCTCC
 751 TTCAACGGTG TGGACTTTGA CATGGTGGGT CGTGGCACCG GCAATCCAAA
 801 TGATGGTTAT GAGGAGTTAA ACCTGAAGTC CACCAAGGGT GACCTCCAGT
 851 TCTCCCCCTG GATTCTGGTC CCTCAAATCG GGTATGGATT CCATCAGTAC
 901 CTGCCCTTCC CCGACGGGAT GTCGCCTTTC CAGGCCGCCA TGAAAGATGG
 951 CTCCGGATAC CAAGTCCATC GCACAATGCA GTTTGAAGAC GGTGCCTCCC
1001 TGACTTCCAA CTACCGCTAC ACCTACGAGG GAAGCCACAT CAAAGGAGAG
1051 TTTCAGGTGA AGGGGACTGG TTTCCCTGCT GACGGTCCTG TGATGACCAA
1101 CTCGCTGACC GCTGCGGACT GGTGCGTGAC CAAGATGCTG TACCCCAACG
1151 ACAAAACCAT CATCAGCACC TTTGACTGGA CTTACACCAC TGGAAATGGC
1201 AAGCGCTACC AGAGCACTGC GCGGACCACC TACACCTTTG CCAAGCCAAT
1251 GGCGGCCAAC ATCCTGAAGA ACCAGCCGAT GTTCGTGTTC CGTAAGACGG
1301 AACTCAAGCA CTCCAAGACC GAACTCAACT TCAAGGAGTG GCAAAAGGCA
1351 TTTACCGATG TGATGTCGAC TGGCACTGGT TCTACGGGCT CGGGCTCCTC
1401 AGGAGAGGAG GATAACATGG CCTCTCTCCC AGCGACACAT GAGTTACACA
1451 TCTTTGGCTC CTTCAACGGT GTGGACTTTG ACATGGTGGG TCGTGGCACC
1501 GGCAATCCAA ATGATGGTTA TGAGGAGTTA AACCTGAAGT CCACCAAGGG
1551 TGACCTCCAG TTCTCCCCCT GGATTCTGGT CCCTCAAATC GGGTATGGCT
1601 TCCATCAGTA CCTGCCCTTC CCCGACGGGA TGTCGCCTTT CCAGGCCGCC
1651 ATGAAAGATG GCTCCGGATA CCAAGTCCAT CGCACAATGC AGTTTGAAGA
1701 CGGTGCCTCC CTGACTTCCA ACTACCGCTA CACCTACGAG GGAAGCCACA
1751 TCAAAGGAGA GTTTCAGGTG AAGGGGACTG GTTTCCCTGC TGACGGTCCT
1801 GTGATGACCA ACTCGCTGAC CGCTGCGGAC TGGTGCGTGA CCAAGATGCT
1851 GTACCCCAAC GACAAAACCA TCATCAGCAC CTTTGACTGG ACTTACACCA
1901 CTGGAAATGG CAAGCGCTAC CAGAGCACTG CGCGGACCAC CTACACCTTT
1951 GCCAAGCCAA TGGCGGCCAA CATCCTGAAG AACCAGCCGA TGTTCGTGTT
2001 CCGTAAGACG GAACTCAAGC ACTCCAAGAC CGAACTCAAC TTCAAGGAGT
2051 GGCAAAAGGC ATTTACCGAT GTGATGGGCA TGGACGAGCT GTACAAG
```

The tandem **Aquamarine-tdLanYFP** was built by replacing the cDNA of Citrine by the one of tdLanYFP in the plasmid coding for the tandem Aquamarine-Citrine between EcoRI and KpnI. [1] The linker between Aquamarine and tdLanYFP was composed of 27 residues and its sequence was SGLRSASVDTMGRDLYDDDDKDPPAEF [1]. It is the same linker for the Aquamarine-Citrine tandem.

**LifeAct-mCitrine and LifeAct-tdLanYFP** were built using the plasmid coding for LifeAct-mTurquoise2 as template (Plasmid #36201, Addgene). The gene coding for mTurquoise2 was replaced by the ones coding for Citrine or tdLanYFP from p-tdLanYFP-N1 and pE-Citrine-N1 plasmids with BamHI and BsrGI restriction enzymes leading to LifeAct-tdLanYFP and LifeAct-Citrine. The mutation A206K was introduced in Citrine by single point mutation and lead to LifeAct-mCitrine (QuikChange site-directed mutagenesis, Stratagene).

For membrane targeting, the shuffled version of the gene coding for tdLanYFP was ordered with BamHI and EcoRI restriction sites at both end of the sequence (Eurofins). They were used to replace directly Dronpa in a Lyn-Dronpa pcDNA3 plasmid (gift from P. Dedecker) leading to **pcDNA3-Lyn-tdLanYFP-*shuffled*** with MGCIKSKRKDKDPQALPVGA as a targeting sequence.

**p-CytERM-mCitrine** and **p-CytERM-tdLanYFP** were built to target the YFPs to the cytosolic side of the endoplasmic reticulum for the **OSER assay**.

The following sequence containing NheI and BsrGI restriction sites (in blue) was ordered (Eurofins):

```
GCAAGCGCTAGCATGGACCCTGTGGTGGTGCTGGGGCTCTGTCTCTCCTGTTTGCTTCTCCTTTCACTCTGGAAACAGAGCTATGGGGGAGGGAAGC
TTCGAATTCTGCAGTCGACGGTACCGCGGGCCCGGGATCCACCGGTCGCCACCATGGTGAGCAAGGGCGAGGAGGATAACATGGCCTCTCTCCCAGC
GACACATGAGTTACACATCTTTGGCTCCTTCAACGGTGTGGACTTTGACATGGTGGGTCGTGGCACCGGCAATCCAAATGATGGTTATGAGGAGTTA
AACCTGAAGTCCACCAAGGGTGACCTCCAGTTCTCCCCCTGGATTCTGGTCCCTCAAATCGGGTATGGCTTCCATCAGTACCTGCCCTTCCCCGACG
GGATGTCGCCTTTCCAGGCCGCCATGAAAGATGGCTCCGGATACCAAGTCCATCGCACAATGCAGTTTGAAGACGGTGCCTCCCTGACTTCCAACTA
CCGCTACACCTACGAGGGAAGCCACATCAAAGGAGAGTTTCAGGTGAAGGGGACTGGTTTCCCTGCTGACGGTCCTGTGATGACCAACTCGCTGACC
GCTGCGGACTGGTGCGTGACCAAGATGCTGTACCCCAACGACAAAACCATCATCAGCACCTTTGACTGGACTTACACCACTGGAAATGGCAAGCGCT
ACCAGAGCACTGCGCGGACCACCTACACCTTTGCCAAGCCAATGGCGGCCAACATCCTGAAGAACCAGCCGATGTTCGTGTTCCGTAAGACGGAGCT
CAAGCACTCCAAGACCGAGCTCAACTTCAAGGAGTGGCAAAAGGCCTTTACCGATGTGATGGGCATGGACGAGCTGTACAACCTGC
```

This sequence codes for CytERM-dLanYFP where dLanYFP is the monomeric version of tdLanYFP. It was cloned in pE-FP-N1 using NheI and BsrGI cloning sites leading to p-CytERM -dLanYFP. The gene coding for dLanYFP was then replaced by the ones coding for tdLanYFP or Citrine using AgeI (in green) and BsrGI restriction sites. The inserts were from p-tdLanYFP-N1 and pE-Citrine-N1 plasmids. Citrine was transformed into mCitrine by introduction of the A206K mutation (QuikChange site-directed mutagenesis, Stratagene).

The plasmid coding **p67-tdLanYFP** was built by replacing the gene of Aquamarine in p-p67-Aquamarine-N1 plasmid using AgeI and BsrGI cloning sites. [3] The plasmid coding for mCherry-p40 was described before[3].

For **AURKA-Aquamarine**, AURKA was subcloned into a pEGFP-N1 vector into the NotI/BamHI cloning sites, while EGFP was replaced by Aquamarine. To obtain **tdLanYFP-AURKA-Aquamarine**, tdLanYFP was inserted into the XhoI/HindIII restriction sites of AURKA-Aquamarine. The Lys162Met variant was obtained from the corresponding wild-type constructs by QuikChange site-directed mutagenesis (Stratagene) with the following primers: 5′-CAAGTTTATTCTGGCTCTTATGGTGTTATTTAAAGCTCAGCT-3′(forward and 5′-AGCTGAGCTTTAAATAACACCATAAGAGCCAGAATAAACTTG-3′(reverse).

### *In vitro* expression and characterization of purified proteins

For **bacterial expression** of dLanYFP, we used commercial plasmid pNCS-dLanYFP (Allele Biotechnology) to produce a 6xHis-tagged protein at its N-terminus. Competent TOP10 cells were transformed with the vector. A volume of 0.5 L of Luria–Bertani (LB) medium containing ampicillin (100 µg.mL$^{-1}$) was inoculated with a 25 mL starter culture that was grown overnight. After 18 h of culture at 30°C, bacteria were harvested by centrifugation (20 min, 5000 rpm) and frozen at -80°C. tdLanYFP, EYFP, Citrine and Venus were expressed as previously described[4]. The proteins were purified with our usual protocol without any modifications[4]. The concentrations of the proteins were measure biochemically with a BCA assay (Bicinchronic acid and copper from Sigma).

**The absorption and fluorescence properties** of purified proteins were studied at 22 °C in a 2 mM HEPES buffer or a 10 mM Tris $H_2SO_4$ buffer pH 8.0 using a Lambda 750 spectrophotometer (Perkin Elmer) and a Fluorolog 3 fluorimeter (Horiba). For absorption, the concentration was typically 10 µM. Solution were diluted more than 20 times for fluorescence monitoring (ODs kept below 0.1). After a vertically polarized excitation at 515 nm (Δλ 2nm), fluorescence emission spectra were collected (Δλ 2nm) with the emission polarizer set at magic angle. After blank subtraction, the fluorescence spectra were corrected according to a transmission curve established using fluorescein in 0.1 M NaOH and rhodamine 6G in EtOH measured with the same set-up[5,6]. Quantum yields were determined at 22°C by preparing a series of parallel dilutions of the protein and fluorescein at ODs ≤ 0.1, and comparing the slope *S* of their integrated corrected spectra as a function of dilution according to:

$$\phi_{FP} = \phi_{Ref} \frac{S_{FP}}{S_{ref}}$$

The reference quantum yield $\Phi_{Ref}$ of fluorescein in 0.1 M NaOH was taken as 0.925 [7].

**The fluorescence lifetimes** of the purified proteins were measured in a 10mM HEPES buffer pH 8 at typical 10 µM concentration and 20 ± 1°C using a TCSPC set-up previously described[8] and a picosecond pulsed excitation (20 MHz repetition rate, P=4µW) from a white light FIANIUM source. The excitation was set at 515 nm (Δλ 5 nm) and fluorescence emission was collected at 535 nm (Δλ 9 nm). The instrumental response function was determined on a LUDOX solution setting the detection wavelength at 515 nm, and was used for reconvolution of analytical models and final comparison to experiments. After best curve fitting, chi-squares were comprised between 1.0 and 1.2 with randomly distributed residuals. In the case of tdLanYFP, the fluorescence lifetimes were obtained by global analysis of decays collected at 535 nm and 580 nm.

**For the determination of $pK_{1/2}$**, the spectra were recorded with a SynergyH1 plate reader (Biotek) in 96-transparent-well plates and spectra were processed as previously described[1]. Absorption spectra were recorded from 240 nm to 600 nm and fluorescence emission spectra from 500 nm to 700 nm upon excitation at 480 nm. For pH levels ranging from 11 to 5.5, buffer solutions contained 33 mM CAPS, 33 mM MES, and 33 mM Bis–tris propane and were adjusted to the appropriate pH by addition of $H_2SO_4$ or NaOH. For pH levels ranging from 5.5 to 2.5, buffer solutions consisted in 50 mM citric acid with appropriate volume of $Na_2HPO_4$ to adjust the pH. Aliquots from a concentrated stock protein solution were diluted into the different buffers at least 12 h before measurements. The concentration of chloride anion [$Cl^-$] was adjusted with a solution of concentrated KCl. Sodium gluconate was added to maintain a constant ionic force of 150 mM (except for the higher chloride concentrations, where no gluconate was added).

**Maturation.** The ability of newly synthetized dLanYFP to mature a fluorescent chromophore was studied in comparison to Citrine as a reference. Top10 bacteria were grown in their usual TB culture medium. In the case of dLanYFP, protein biosynthesis and fluorescence expression took place constitutively all along bacterial growth in LB medium. In the case of Citrine, protein expression in TOP10 cells were induced after prior cell growth to approximately 1 OD, by adding IPTG and ampicillin. Samples of the bacterial culture were collected at different times (typically every 30 min from 1 h to 5 h) after culture seeding (dLanYFP) or induction of protein expression (Citrine), centrifuged (20 min, 5000 rpm) and the cell pellets were immediately stored frozen at -80°C. On the day of experiments, pellets were thawed and cells were incubated under gentle agitation for 15 min in 2 mL of a lytic buffer (Lyse Cell Lytic B, Sigma in 40mM Tris-HCl pH8.0) containing 2mM PMSF as a protease inhibitor, followed by 2 min centrifugation at 15000 rpm. The progressive increase in fluorescence in the supernatant was then monitored using a SynergyH1 plate reader (Biotek). Kinetic traces followed a similar time course independent of the sampling time of the bacterial culture for dLanYFP, while in the case of Citrine, the apparent fluorescence growth was progressively slower with increasing sampling time, which we ascribed to different levels of concurrent protease activity. Accordingly, only bacterial samples collected during the first two hours after induction were used. Averaged experimental results are presented Figure S7.

Supplementary Figures

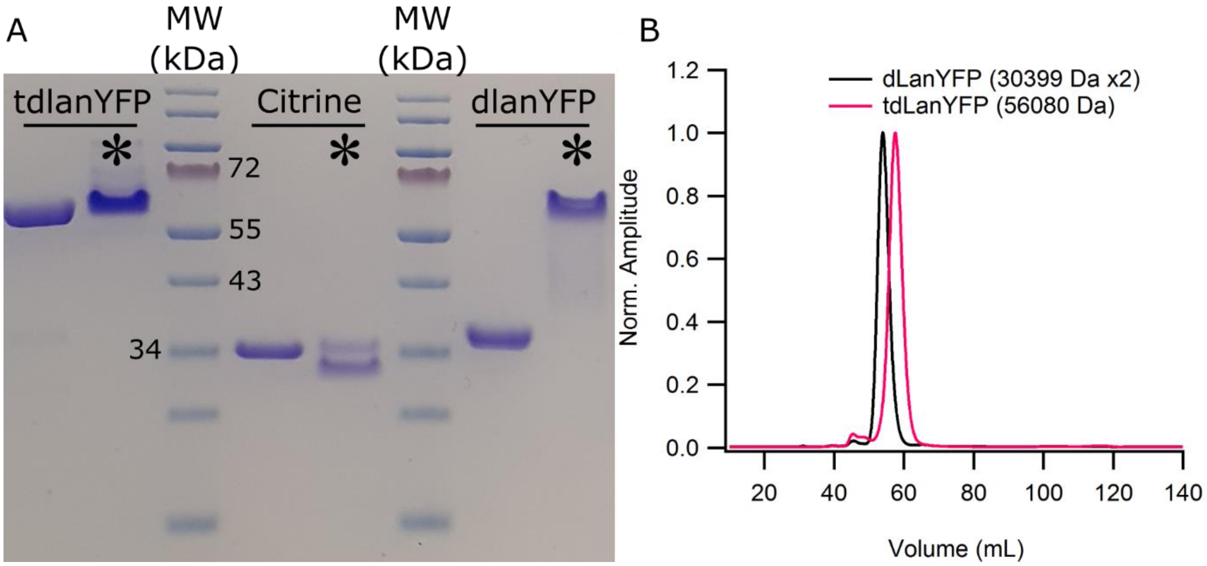


**Figure S1:** (A) SDS-PAGE of purified tdLanYFP, Citrine and dLanYFP. In lanes with a star, the samples were not heated whereas the samples in lanes 1, 4 and, 7 were heated 10 min at 90°C, Size Marker EZ-RUN Pre-Stained *Rec* Protein Ladder. (B) Chromatograms from the gel filtration for dLanYFP and tdLanYFP (Column HiLoad 16/600, Superdex 75pg GE Healthcare Life Sciences). More than 95% of the proteins are in the main peak. The intermolecular dimer of dLanYFP (MW 60598 Da) has an elution volume slightly smaller than tdLanYFP consistently with their relative molecular weight.

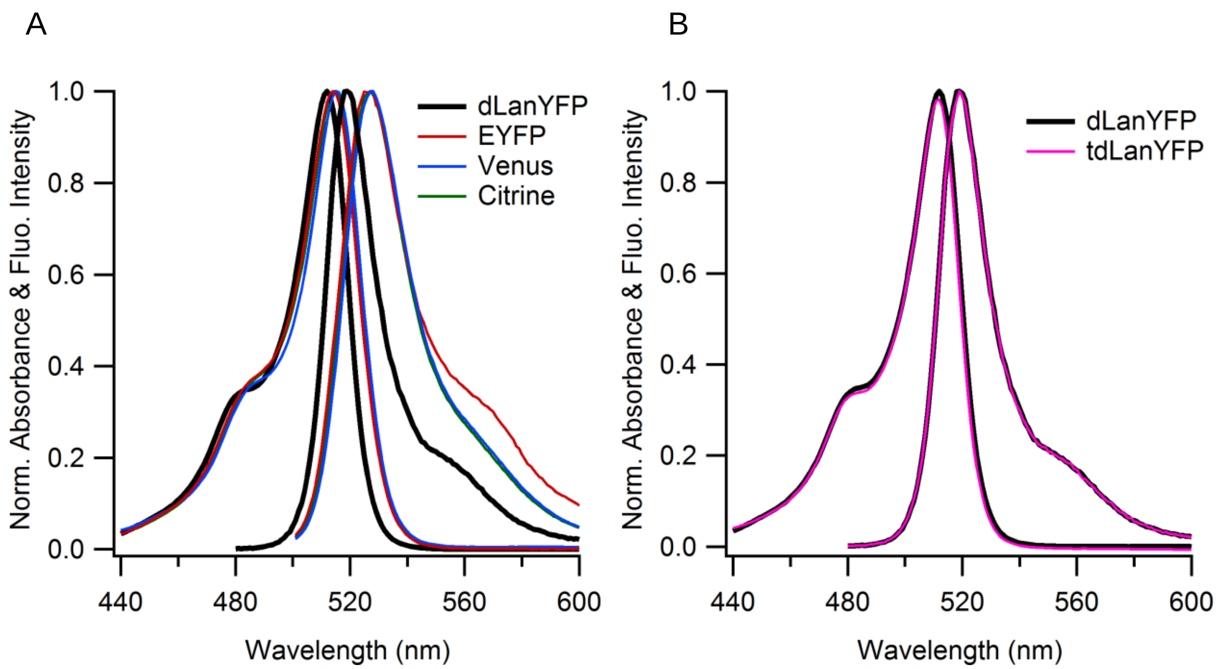


Figure S2: Normalized absorption and emission spectra of dLanYFP, EYFP, Venus and Citrine A). Comparison of dLanYFP and tdLanYFP (B). The quantum yields of dLanYFP and tdLanYFP differ from 2% (within the experimental uncertainty of 5%). Emission spectra were recorded with excitation at 515 nm.

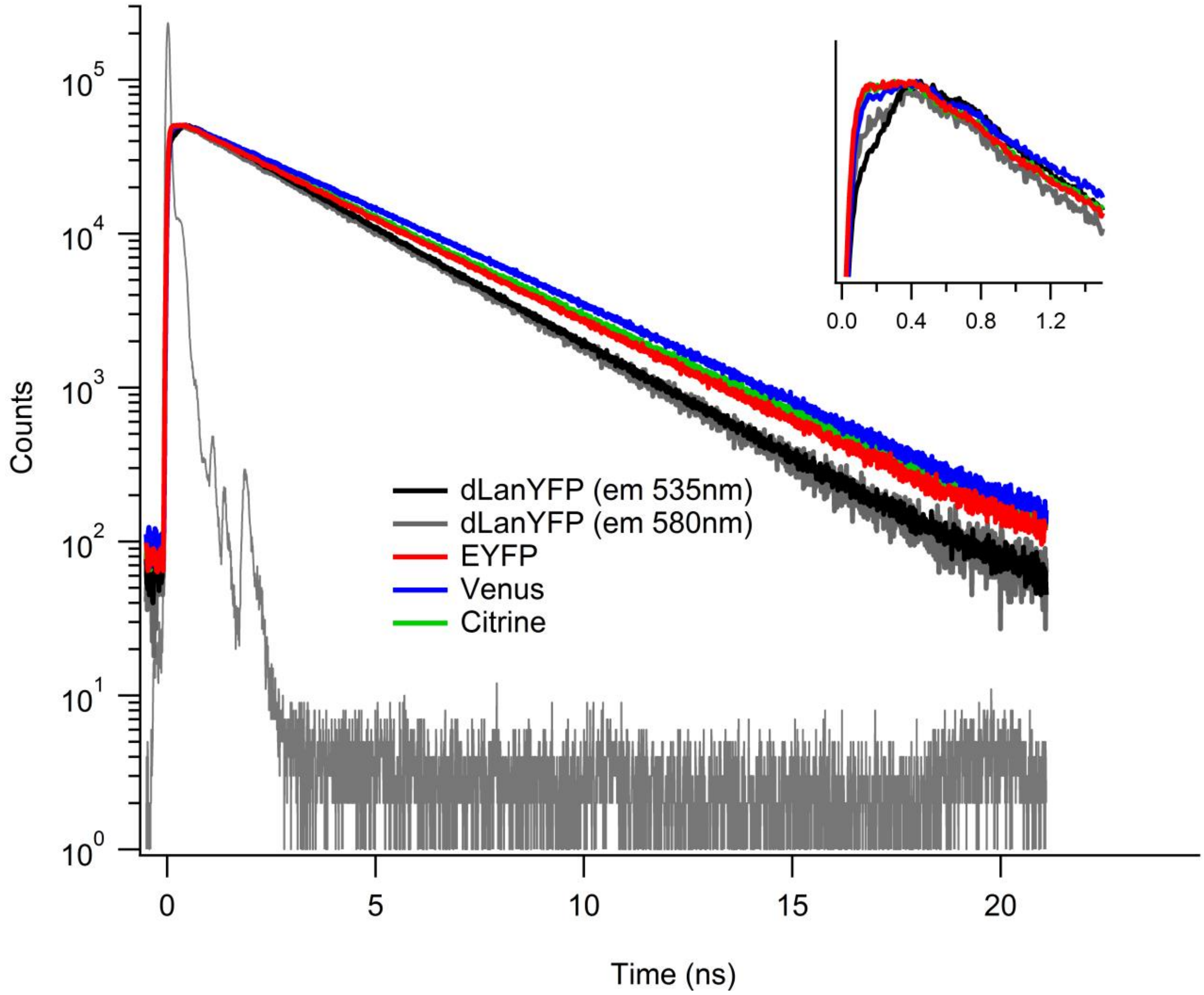


**Figure S3**: Fluorescence decays of purified yellow fluorescent proteins acquired upon excitation at 515 nm and detection at 535 nm ($\Delta\lambda$=9 nm), otherwise it is specified. The instrumental response function was determined on a LUDOX solution setting the detection wavelength at 515 nm. Inset: Zoom on the region between 0 and 1.5 ns

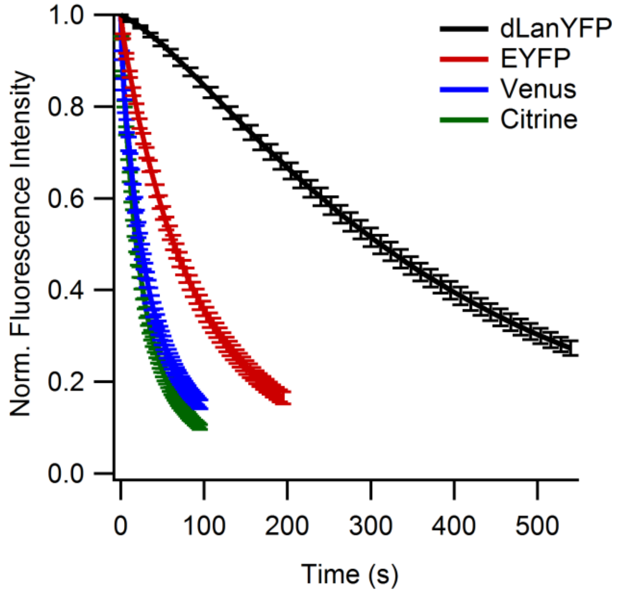


**Figure S4:** Irreversible photobleaching of YFPs under continuous irradiation in wide-field illumination (0.4 W/cm²). FPs are fixed on $Ni^{2+}$-NTA agarose beads through their His-tag. Solid lines correspond to the best fits to an exponential analytical model.

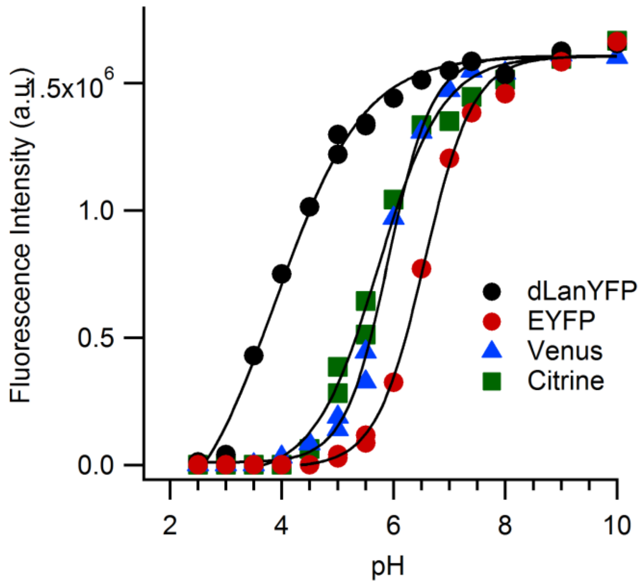


**Figure S5:** pH dependence of the fluorescence intensity of proteins excited at 480 nm and detected at their maximal emission. Solid lines correspond to the best fits to a sigmoidal analytical model. Experimental data were normalized to 100% maximum of their analytical fits.

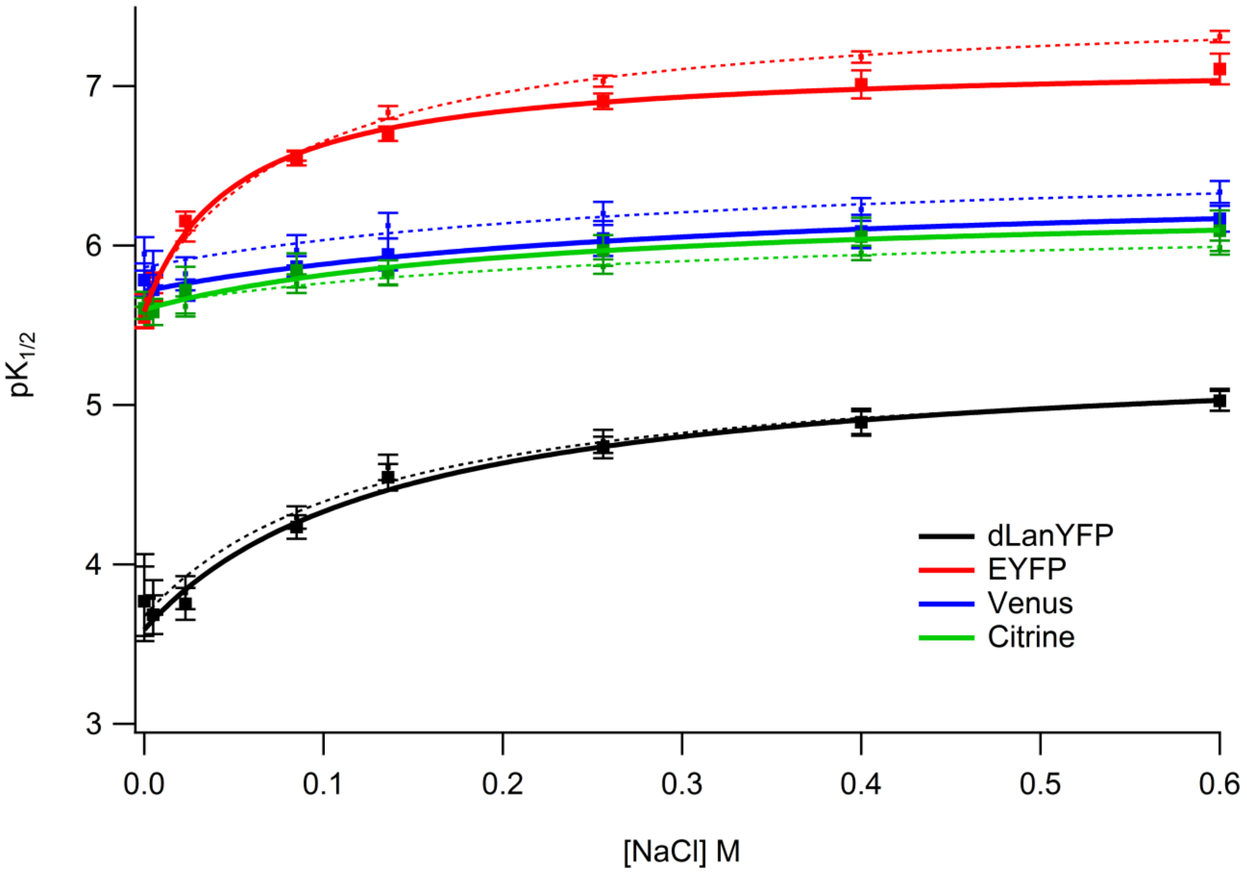


**Figure S6**: Chromophore protonation $pK_{1/2}$ of purified yellow fluorescent proteins as a function of chloride concentration. The chromophore $pK_{1/2}$ were obtained from studies of their absorption spectra (squares) or of their fluorescence emission spectra (dots) as a function of pH performed at varying NaCl concentrations. Lines represent best fits of the data to a saturation hyperbole (absorption, continuous lines and fluorescence, dashed lines): $pK_{1/2}[NaCl] = \frac{pK_{max}[NaCl]+pK_{min}\times K_{app}}{[NaCl]+K_{app}}$.

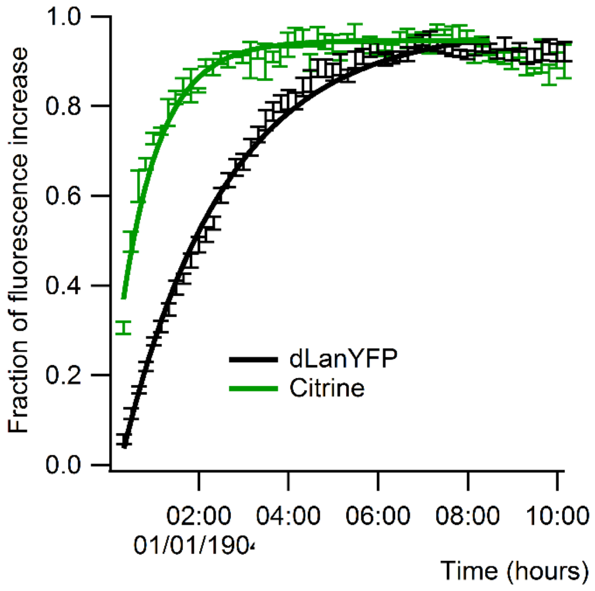


**Figure S7**: Comparison of maturation rates of dLanYFP and Citrine expressed in *E. coli*.

MVSKGEEDNMA SLPATHELHIFGSFNGVDFDMVGRGTGNPNDGYEELNLK 50
STKGDLQFSPWILVPQIGYGFHQYLPFPDGMSPFQAAMKDGSGYQVHRTM 100
QFEDGASLTSNYRYTYEGSHIKGEFQVKGTGFPADGPVMTNSLTAADWCV 150
TKMLYPNDKTIISTFDWTYTTGNGKRYQSTARTTYTFAKPMAANILKNQP 200
MFVFRKTELKHSKTELNFKEWQKAFTDVMSTGTGSTGSGSSGEEDNMASL 250
PATHELHIFGSFNGVDFDMVGRGTGNPNDGYEELNLKSTKGDLQFSPWIL 300
VPQIGYGFHQYLPFPDGMSPFQAAMKDGSGYQVHRTMQFEDGASLTSNYR 350
YTYEGSHIKGEFQVKGTGFPADGPVMTNSLTAADWCVTKMLYPNDKTIIS 400
TFDWTYTTGNGKRYQSTARTTYTFAKPMAANILKNQPMFVFRKTELKHSK 450
TELNFKEWQKAFTDVM GMDELYK

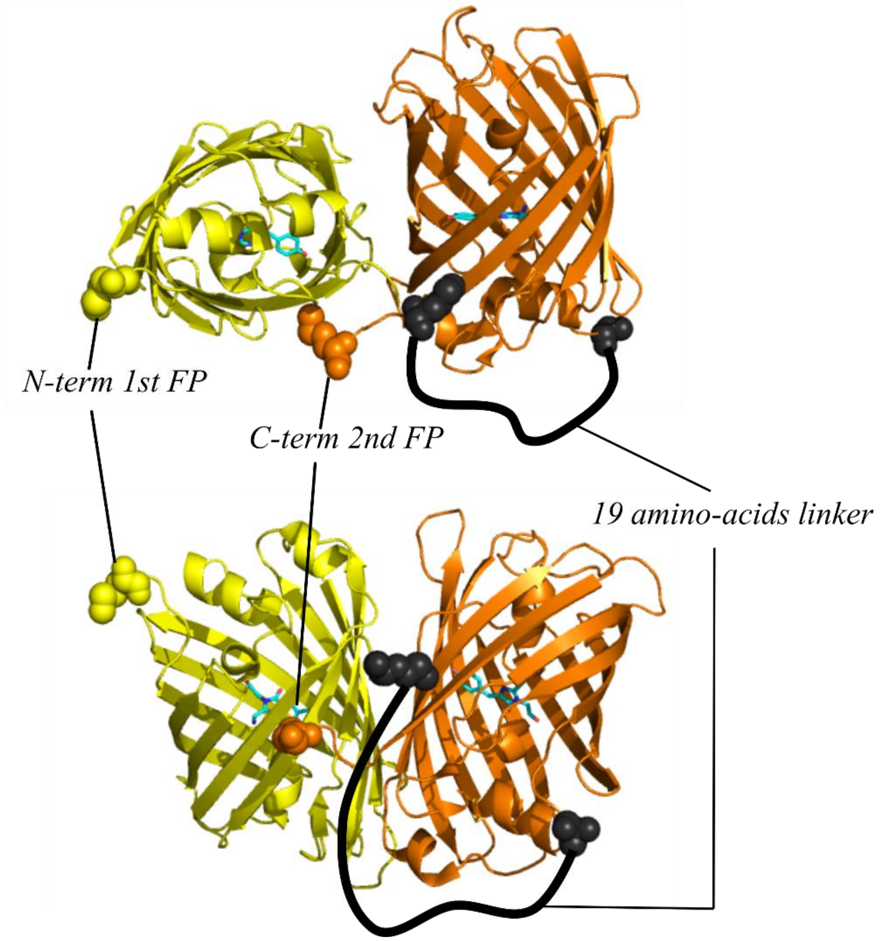


**Figure S8**: Amino-acid sequence and schematic structure of the tandem tdLanYFP. The amino-acid sequence of each dLanYFP is emphasized in yellow and orange, while the linker is in black. In the structure, the N and C termini of each FP are in ball representation and the linker between the two dLanFP is the black line. The scheme of the tandem was built using the tetramer structure, pdb file 5LTQ [10]. The working name of tdLanYFP was superYFP [11].

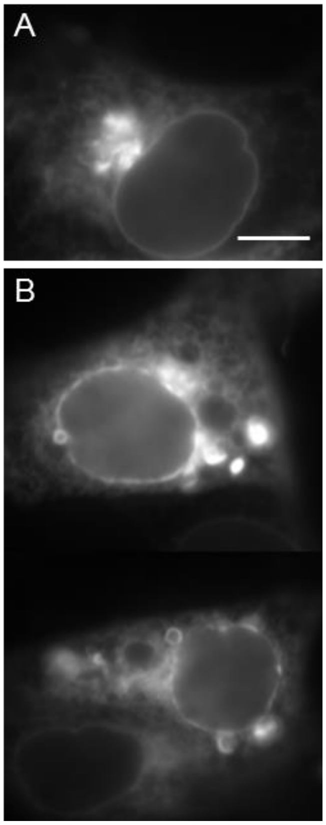


**Figure S9**: Wide-field fluorescence images of representative OSER positive cells. CytERM-mCitrine (A) and CytERM-tdLanYFP (B) fusion proteins were expressed in COS7 cells. Scale bar 10 µm.

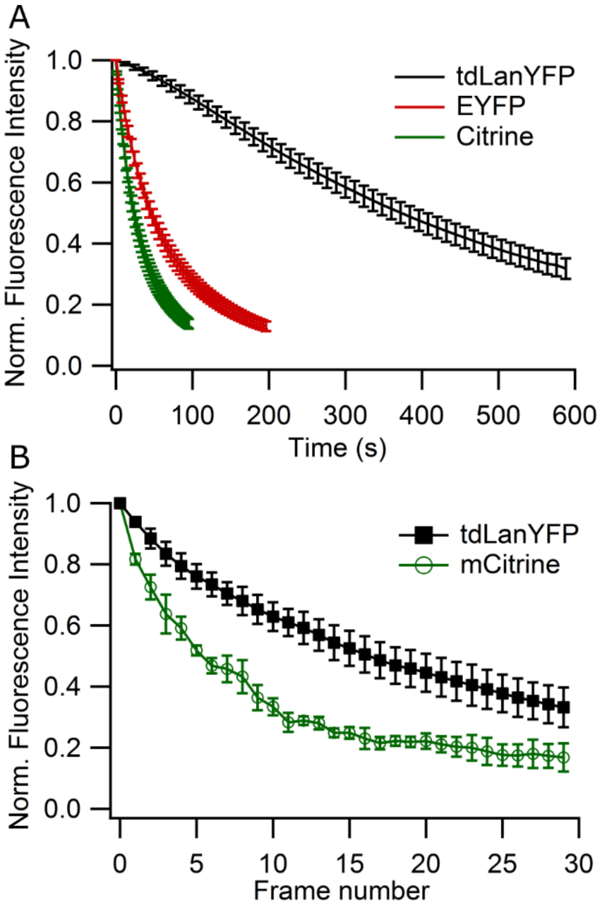


**Figure S10:** Irreversible photobleaching of YFPs in live COS7 cells. (A) Decay of the fluorescence intensity in COS7 cells expressing cytosolic tdLanYFP, EYFP or Citrine under continuous irradiation in wide-field illumination (0.4 W/cm²). Solid lines correspond to the best fits to an exponential analytical model. Each point is the average of 5 to 10 cells. (B) Decay of the fluorescence intensity of COS7 cells expressing LifeAct-tdLanYFP or LifeAct-mCitrine during a 30-frame time-lapse acquired by confocal microscopy (Average power measured during the scanning on the objective 1,2 µW). Each point is the average of 5 ROIs.

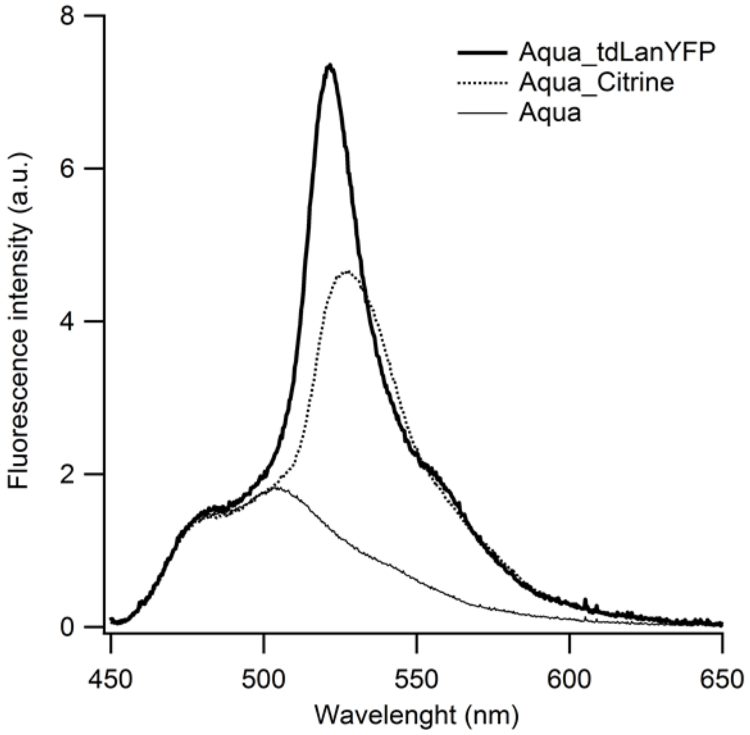


**Figure S11**: Fluorescence spectra normalized at 480 nm of a whole field of view of COS7 cells transfected with the Aquamarine donor alone or with the FRET fusion of Aquamarine-acceptor, Citrine or tdLanYFP (excitation wavelength at 438 nm with the CFP cube without emission filter, see Table S1).

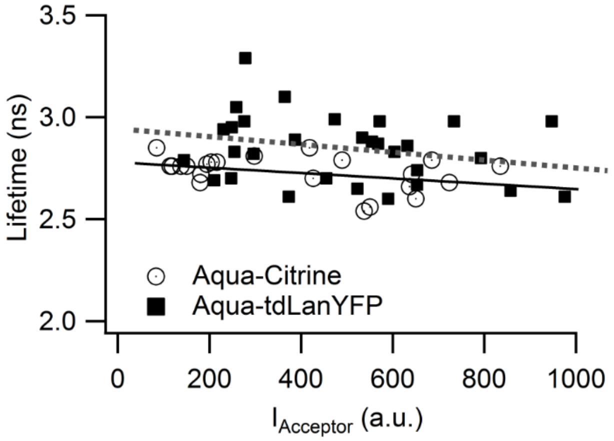


**Figure S12**: Correlation between the donor lifetime and the fluorescence intensity of the acceptor, $I_{Acceptor}$, for tandem expressed in COS7 cells. We set a maximum limit for $I_{Acceptor}$ at which the donor lifetime decreased less than 5 % ($I_{Acceptor}$ < 1000 a.u.) to limit the so-called molecular crowding.

Supplementary Tables

| cube | excitation filter | dichroic mirror | emission filter |
|---|---|---|---|
| CFP (CFP 2432C) [a] | FF02-438/24-25 | FF458-Di02 | FF01-483/32-35 |
| YFP (YFP 2427B) [a] | FF01-500/24-25 | FF520-Di02 | FF01-542/27-25 |
| RFP (C156423 custom) [b] | ET580/25x | ZT594rdc | ET625/30m |

([a] Semrock, Rochester, NY, [b] Chroma Technology Corp., Bellows Falls, VT).

**Table S1**: The spectral selections for CFP, YFP, FRET and RFP in the epifluorescence pathway of the TCSPC-FLIM microscope

| Excitation wavelength | Filter1 | Filter2 | Filter3 |
|---|---|---|---|
| 440nm | > 458 nm | > 458 nm | 480 ± 15 nm |
| 466 nm | > 488 nm | | 535 ± 20 nm |

**Table S2**: The spectral selections for CFP and YFP fluorescence emission before the MCP-PMT detector in the TCSPC pathway of the TCSPC-FLIM microscope.
